# Effects of N-linked glycan of Lassa Virus Envelope Glycoprotein on the Immune Response

**DOI:** 10.1101/2020.09.29.319855

**Authors:** Xueqin Zhu, Yang Liu, Jiao Guo, Zonglin Wang, Junyuan Cao, Gengfu Xiao, Wei Wang

## Abstract

Lassa virus (LASV) belongs to the *Mammarenavirus* genus (family Arenaviridae) and causes severe hemorrhagic fever in humans. The glycoprotein precursor (GPC) contains eleven N-linked glycans that play essential roles in GPC functionalities such as cleavage, transport, receptor recognition, epitope shielding, and immune response. We used three mutagenesis strategies to abolish the individual glycan chains on the GPC and found that all three mutations led to cleavage inefficiency on the 2^nd^, 5^th^, and 8^th^ glycosylation motifs. To evaluate N to Q mutagenesis for further research, it was found that deletion of the 2^nd^ and 8^th^ glycans completely inhibited the infectivity. We further investigated the role of glycans on GPC-mediated immune response by DNA immunization of mice. Deletion of the individual 1^st^, 3^rd^, 5^th^ and 6^th^ glycans significantly enhanced the proportion of effector CD4+ cells, whereas deletion of the 1^st^, 2^nd^, 3^rd^, 4^th^ 5^th^, 6^th^, and 9^th^ glycans enhanced the proportion of CD8+ effector T cells. Deletion of specific glycans improves the Th1-type immune response, and abolishment of glycan on GPC generally increases the antibody titer to the glycan-deficient GPC. However, the antibodies from either the mutant or WT GPC-immunized mice show little neutralization effect on wild-type LASV. The glycan residues on GPC provide an immune shield for the virus, and thus represent a target for the design and development of a vaccine.

**Importance:** At present, there are no Food and Drug Administration-approved drugs or vaccines specific for LASV. Similar to other enveloped viruses with a heavy glycan shield, the N-linked glycans of LASV make it difficult for effector T cells and neutralization antibodies to access the glycoprotein epitope. In this study, we evaluated the effect of the individual glycan chains on GPC-mediated immune response, and found that deletion of the glycan improves the proportion of effector T cells, improving the Th1-type immune response, and increasing the antibody titer to the WT and mutant GPC, which may be beneficial to vaccine design and development.

## Introduction

Lassa virus (LASV) belongs to genus *Mammarenavirus*, family *Bunyaviridae*. The natural reservoir of LASV is *Mastomys natalensis* in Africa, and humans are infected through direct contact with their excreta or exposure to the aerosol. Between 300,000 and 500,000 people are infected with LASV annually, and the mortality of hospitalized patients ranges from 20% to 70%. The United States Centers for Disease Control and Prevention classifies the virus as a Category A bioterrorism agent, and there is currently no drug or vaccine approved by the Food and Drug Administration capable of treating or preventing Lassa fever.

The envelope glycoprotein complex (GPC) of LASV is sequentially cleaved by signal peptidases and subtilisin kexin isozyme-1 (SKI-1)/site-1 protease (S1P) enzyme during the maturation process to obtain the stable signal peptide (SSP), receptor-binding GP1, and envelope fusion protein GP2. *Mammarenavirus* GPC is a heavily glycosylated protein. It has been estimated that N-linked glycosylation accounts for nearly 30% of the total mass of LASV GPC (1–3). The LASV lineage IV Josiah strain is the most commonly used strain in the development of LASV vaccines. It has 11 N-glycosylation motifs (Asn-X-Thr/Ser, where X is any amino acid except proline) on its GPC (Fig. 1A). These 11 N-linked glycans are distributed relatively evenly on the surface of the GPC in terms of spatial conformation, with seven glycans on GP1 and four on GP2. Glycans play critical roles in many biological functions associated with the GPC, such as cleavage, folding, receptor recognition, epitope shielding, and immune response (4). Investigation on hospitalized patients infected with LASV found that a small amount of neutralizing antibodies could be detected in only a few convalescents at a relatively late stage of disease course, and that the neutralizing power of these antibodies was relatively low (4–8). However, a previous study found that the cellular immune response plays a major role in immune defense against the LASV (9, 10). We hypothesized that in LASV, the large number of N-linked glycans on the surface of the virion shields important GPC epitopes, which adversely affects immune recognition of the virus and inhibits induction of a specific immune response, thereby making treatment of and rehabilitation from Lassa fever difficult. Therefore, studying the role of the 11 N-sugar chains on the GPC of LASV with regards to the host immune response will help identify the mechanisms of immune escape employed by the virus and facilitate subsequent development of effective vaccines or therapeutic antibodies.

**Fig. 1.**
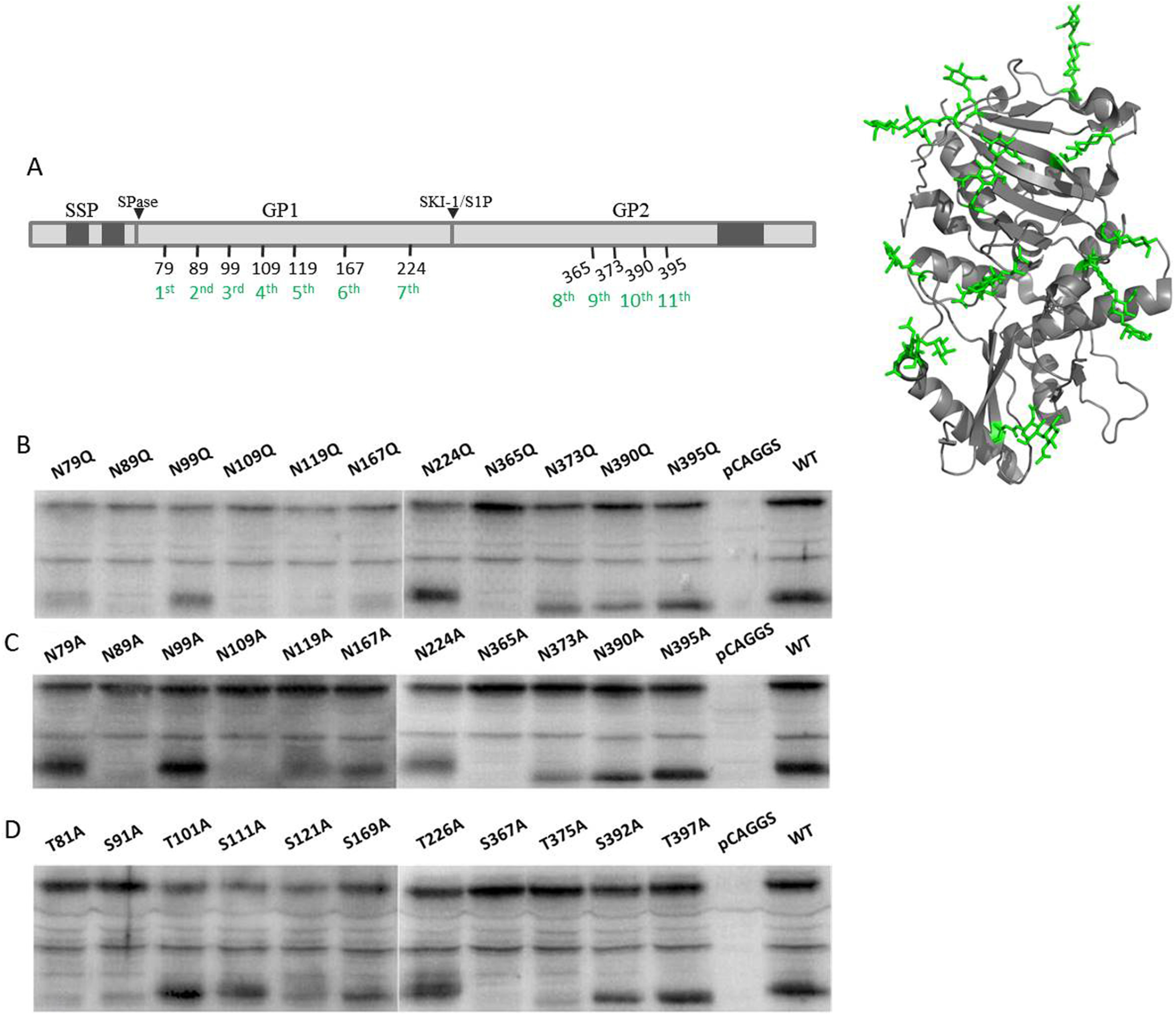
Proteolytic cleavage of the individual N-linked glycosylation motif mutants. (A) Schematic diagram and cartoon representation (PDB: 5VK2 (1)) of the glycosylation of LASV GPC. The precursor glycoprotein GPC was cleaved by SPase and SKI-1/S1P. The mature GPC contains SSP (1–58), GP1 (59–259), and GP2 (260–491) domains and is modified by 11 N-linked glycan chains. Transmembrane domains are indicated in gray. (B - D) HEK 293T cells expressing the wild-type GPC and individual N-glycosylation mutants of N→Q (B), N→A (C), and S/T→A (D), respectively. The expressed proteins were separated via SDS-PAGE and western blotting was carried out using anti-GP2 antisera. Images chosen are representative of at least three independent assays.

## Results

### Effects of N-glycosylation Modification on GPC Cleavage

To investigate the influences of different mutations of the 11 glycosylation motifs upon GPC cleavage, the corresponding asparagine residues were individually mutated to glutamine or alanine, and the serine/tyrosine residues in the motif were mutated to alanine. Thus, 33 mutations were constructed, and western blotting was performed using antiserum against GP2. As shown in Fig. 1, all mutations of the N-glycan motifs on GP2 resulted in slight decreases (~2 kDa) in the molecular weight of the respective GP2 bands relative to the wild-type (WT) GP2 band.

Next, we determined the effect of the N-linked glycans on protease cleavage by testing for the presence of cleaved GP2. Disruption of the N-linked glycosylation motif by substitution of asparagine with the structurally similar glutamine on the 2^nd^, 4^th^, 5^th^, and 8^th^ motifs (corresponding to N89Q, N109Q, N119Q, and N365Q) inhibited proteolytical processing, and those with deletions at the remaining seven motifs (corresponding to N79Q, N99Q, N167Q, N224Q, N373Q, N390Q, and N395Q) were unaffected (Fig. 1B). Similarly, disruption of the glycosylation motifs by substitution of asparagine with alanine on the 2^nd^, 4^th^, and 8^th^ motifs (corresponding to N89A, N109A, and N365A) abolished proteolytical processing, and the mutations in the 5^th^ and 9^th^ (corresponding to N119A and N373A) exerted mild inhibition, whereas the other N to A mutations had no influence (Fig. 1C). Intriguingly, substitution of serine or threonine at the 8^th^ glycosylation site with alanine (corresponding to S367A) abolished the GP1-GP2 cleavage, and the mutations in the 1^st^, 2^nd^, 5^th^, and 9^th^ (corresponding to T81A, S91A, S121A, and T375A) exerted mild inhibition on the cleavage efficiency (Fig. 1D). It was shown that disrupting the glycosylation motifs by introducing different mutations led to differing results, which might be due to the changes of the residue per se, rather than the loss of the specific glycan. However, all three mutations led to a decrease in cleavage efficiency on the 2^nd^, 5^th^, and 8^th^ glycosylation motifs, suggesting that the 2^nd^, 5^th^, and 8^th^ N-linked glycans were indispensable for GP1-GP2 cleavage. Given the consistency in the results for the N to A and N to Q substitutions, and that the structures of N and Q are the most similar, we used the N to Q mutation for further research, which caused a minimal change to the spatial structure of LASV GPC while ensuring glycosylation-site mutation.

### Effects of N-glycosylation Modification on the infectivity of LASV Psuedotype Virus

To evaluate the influence of the individual glycan on the pseudotype virus infectivity, we constructed the pseudotype viruses with the VSV backbone bearing the mutant GPC. The infection activities were evaluated in Vero cells (Fig. 2). Disruption of the N-linked glycans by the N to Q substitutions on the 2^nd^, 4^th^, 5^th^, and 8^th^ glycosylation motifs led to a significant loss of LASVpv (LASV pseudovirus) infectivity when compared with LASVpv packaged with WT GPC. Deletion of the 2^nd^ and 8^th^ N-linked glycans completely inhibited the infectivity, and deletion of the 4^th^ and 5^th^ N-linked glycans led to partial inhibition. These results were in line with the protease cleavage results depicted above, indicating that efficient cleavage of premature GPC was a prerequisite for downstream function.

**Fig. 2.**
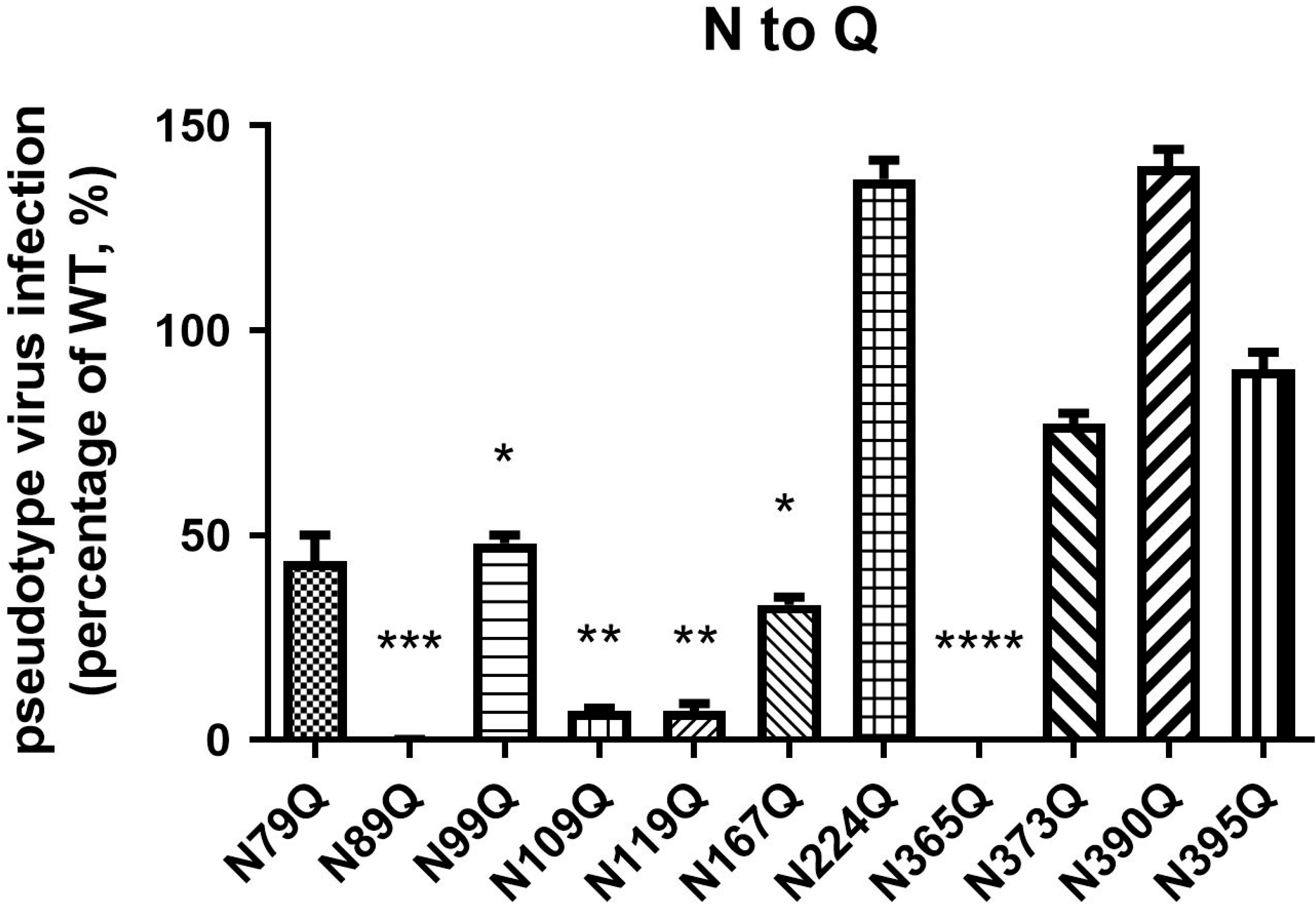
The infectivities of pseudotype viruses bearing mutant GPC. The genome copies of the pseudotype viruses bearing the WT and mutant GPC were quantified using qPCR. Vero cells were infected with the WT and mutant viruses with the same genome copies and the Rluc were determined 24 hours later. Data are presented as mean ± SD of at least 3 independent assays. *P < 0.05, **P < 0.01, compared to WT.

### Effects of N-glycosylation Modification on Effector CD4+ T Cells and CD8+ T Cells among Spleen Lymphocytes

The 11 N-linked glycans on LASV GP play an important role in GPC cleavage and maturation, as well as in pseudovirus infection. Additionally, N-glycosylation modification is involved in various aspects of the immune response to GPC, such as receptor recognition on the host cell surface protecting LASV from the immune response (3, 4, 11–13).

Therefore, we focused on how modification of the 11 N-linked glycans on LASV GP influences the immune response to LASV GPC. We employed recombinant GPC plasmids with N to Q substitution as the DNA vaccine used to immunize BALB/c mice. BALB/c mice were immunized with 13 groups (6 mice per group) of DNA vaccines, including 11 groups of mutated recombinant plasmids, pCAGGS-GPC_N → Q_, WT GPC, and control pCAGGS. Immunofluorescence analyses of CD3, CD4, CD8, and interferon (IFN)-γ were performed (Fig. 3). The proportions of CD3+ T cells in spleen lymphocytes in 78 BALB/c mice across the 13 vaccinated groups were within the range (~40%) of those observed during normal immune response, whereas the deletion of the 1^st^, 2^nd^, 4^th^, and 8^th^ (N79Q, N89Q, N109Q, N365Q) glycans caused the proportion of CD3+ T cells to decrease slightly when compared to the WT (Fig. 3A). Moreover, abolishment of the 1^st^, 2^nd^, and 6^th^ (N79Q, N89Q, N167Q) glycans caused the proportion of CD4+ T cells to decrease, whereas abolishment of the 5^th^, 9^th^, and 10^th^ (N119Q, N373Q, N390Q) glycans led to the proportion of CD8+ T cells to increase (Fig. 3B and 3C).

**Fig. 3.**
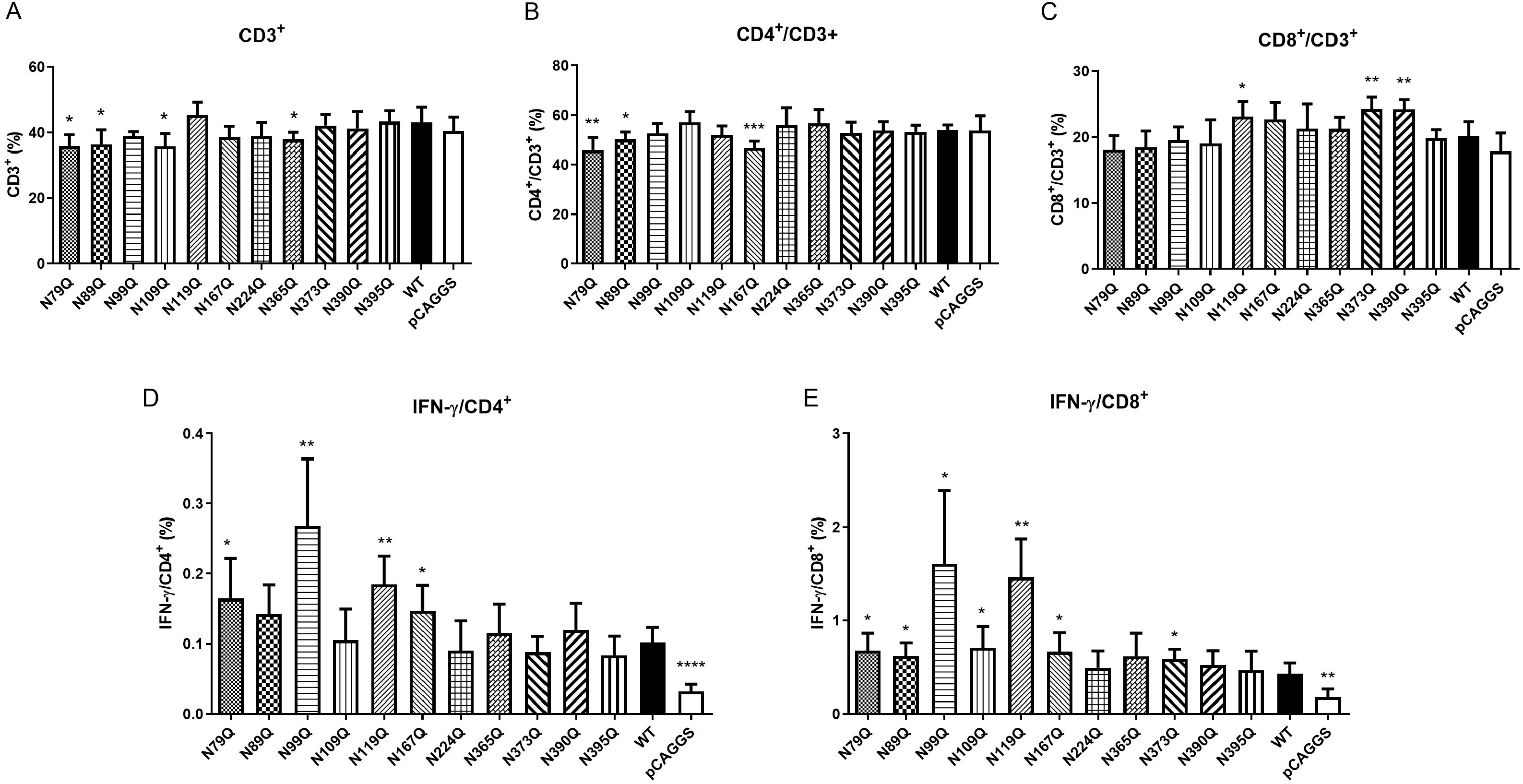
Effects of N-glycan deletion on T cells in spleen lymphocytes. The mouse spleen lymphocytes were separated, cultured, and stained with phycoerythrin (PE)-conjugated rat anti-mouse CD8a, fluorescein isothiocyanate-conjugated rat anti-mouse CD4, PE– CyTM7-conjugated hamster anti-mouse CD3e, and 7-AAD viability staining solution (BioLegend) at 4°C for 20 min in the dark. (A–C) Effects of individual N-glycosylation deletions on CD3+ (A), CD4+ (B), and CD8+ (C) T cells in spleen lymphocytes. Effector T cells CD4+ (D) and CD8+ (E) T cells were stained with allophycocyanin-conjugated rat anti-mouse IFN-y antibody (1:200). Each group contained five - six mice. *P < 0.05, **P < 0.01, ***P < 0.001, compared to WT.

Lymphocytes that secrete IFN-γ represent an effector subset of these cells. First, we found that immunization with the WT resulted in a significant increase in the proportion of both the effector CD4+ and CD8+ cells relative to the control plasmid vaccine group. Moreover, deletion of the 1^st^, 3^rd^, 5^th^ and 6^th^ (N79Q, N99Q, N119Q, N167Q) N-linked glycans on GPC significantly enhanced the proportion of effector CD4+ cells, and deletion of the remaining seven N-sugar chains had no effect on their proportion. For CD8+ effector T cells, deletion of the 1^st^, 2^nd^, 3^rd^, 4^th^ 5^th^, 6^th^, and 9^th^ N-linked glycans enhanced their proportion relative to IFN-γ, and deletion of the 7^th^, 8^th^, 10^th^, and 11^th^ N-linked glycans had no effect (Fig. 3D and E).

### Effects of N-linked Glycans on Cytokines Secreted by Spleen Lymphocytes

Among the seven cytokines tested, interleukin (IL)-2, IFN-γ, and tumor necrosis factor (TNF)-α are associated with the Th1-type cellular immune response, IL-4, IL-6, and IL-10 with the Th2-type response, and IL-17A to the Th17-type response. Fig. 4 shows that all seven cytokines, except for IL-17A, increased in WT group relative to control pCAGGS group. Specifically, deletion of the individual 3^rd^ to 10^th^ N-linked glycans on GPC significantly increased IL-2 levels in spleen lymphocytes relative to those in mice receiving WT GPC (Fig. 4A), whereas deletion of the individual 3^rd^, 5^th^, 6^th^, 8^th^, 9^th^, and 10^th^ glycan increased IFN-γ levels (Fig. 4B); TNF-α was only slightly elevated when either the 3^rd^ or 9^th^ glycans were abolished (Fig. 4C).

**Fig. 4.**
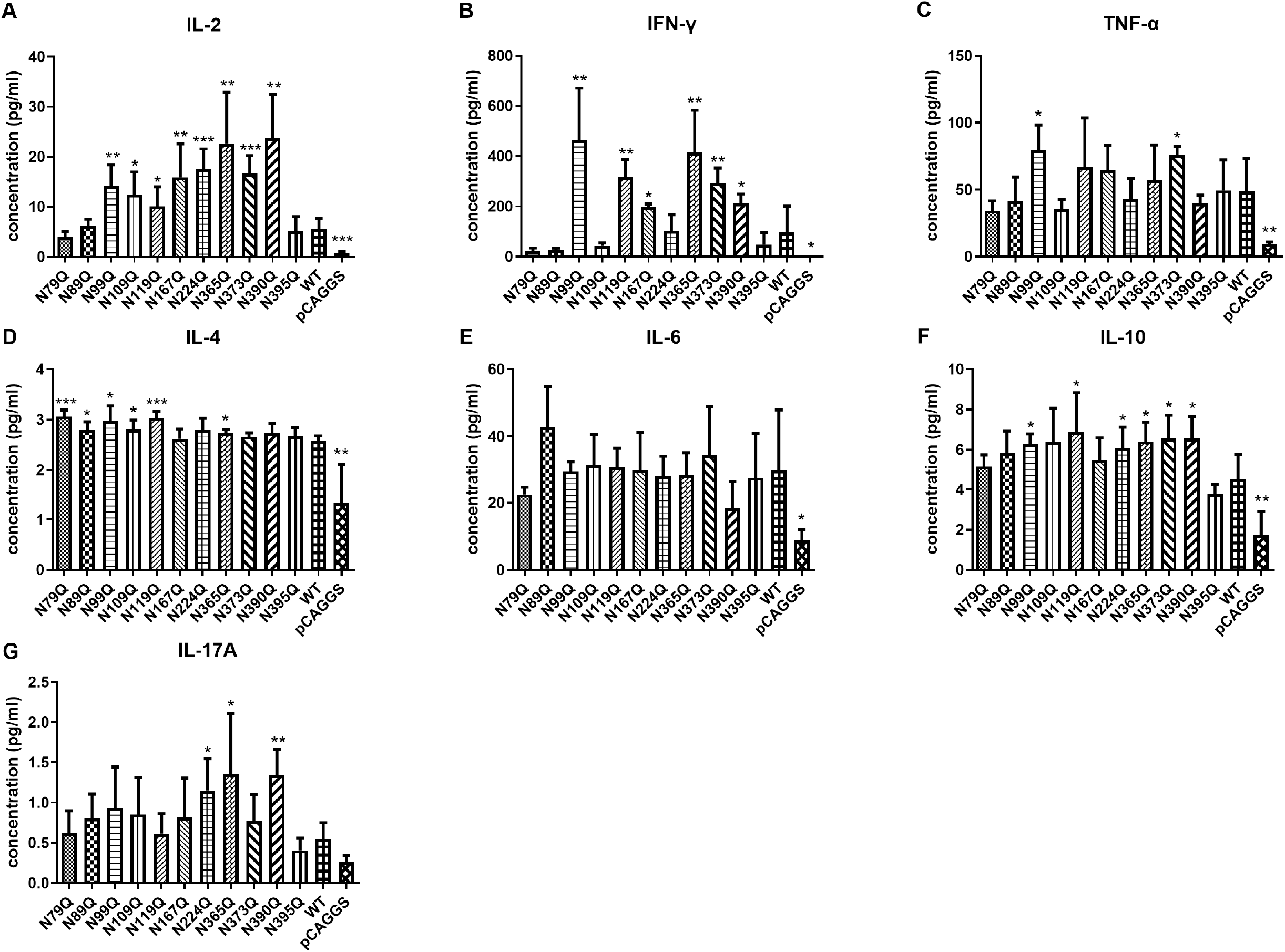
Effects of N-glycan deletions on cytokines secreted by spleen lymphocytes were labeled by using a cytometric bead array kit (BD Biosciences), and IL-2 (A), IFN-γ (B), TNF-α (C), IL-4 (D), IL-6 (E), IL-10 (F), and IL-17A (G) were determined using flow cytometry. Each group contained five - six mice. *P < 0.05, **P < 0.01, *** P < 0.001, compared to WT.

Furthermore, we found that deletion of the 1^st^ and 5^th^ N-linked glycans on GPC significantly increased IL-4 secretion, deletion of the 2^nd^, 3^rd^, 4^th^, and 8^th^ slightly increased IL-4 secretion, and deletion of the remaining five glycans had no effect (Fig. 4D). However, IL-6 levels in either glycan mutation motif showed no significant difference relative to levels in the WT group (Fig. 4E). Furthermore, deletion of the 3^rd^, 5^th^, 7^th^, 8^th^, 9^th^, and 10^th^ N-linked glycans slightly increased IL-10 levels, whereas deletion of the remaining five N-linked glycans had no effect (Fig. 4F).

When studying induction of the Th17-type immune response (Fig. 4G), we observed elevated levels of IL-17A in spleen lymphocytes from mice in the 7^th^, 8^th^, and 10^th^ deletion mutation groups relative to those in mice receiving the WT vaccine.

These results indicated that deletion of N-linked glycans on GPC, especially those at the 3^rd^, 5^th^, 8^th^, and 9^th^ sites, primarily enhanced the induction of both Th1 and Th2 immune response. Intriguingly, deletion of the 11th glycan on GPC had no effect on Th1, Th2, and Th17A immune response. These findings suggest these specific glycan chains assist LASV in escaping immune response by reducing host recognition of GPC and precluding induction of the immune response.

### Effects of N-glycosylation Modification on Antibody Titers

We then detected antibody titers obtained from the individual glycan deletion mutation immunized serum against each mutated GPC variant using cell-based ELISA. Fig. 5 shows that antibody titers in mice receiving mutated GPC variants as well as the WT GPC plasmid generally had a higher affinity for the glycan deletion mutant GPC than the WT GPC, suggesting that the glycans on LASV GPC might play a role in immune escape by shielding the epitope and thus making it inaccessible to antibodies. Evaluation of which N-glycosylation site affected antibody titers revealed that deletion of N-linked glycan at the 3^rd^, 5^th^, 6^th^, 8^th^, and 9^th^ sites significantly increased the antibody titer. The titers obtained from these five groups could reach to approximately 17(-log2) whereas the titers from the remaining seven groups were approximately 15(-log2) (Fig 5A-L). Similarly, to compare the antibody titers when titrated with WT GPC, we found that deletion of the 3^rd^, 5^th^, 6^th^, 8^th^, and 9^th^ N-linked glycans generated significantly higher antibody titers than other deletion mutations as well as WT (Fig. 5M), thereby indicating a role for the 3^rd^, 5^th^, 6^th^, 8^th^, and 9^th^ N-glycosylation sites in the GPC in immune escape and reduced humoral immune response by shielding key GPC epitopes.

**Fig. 5.**
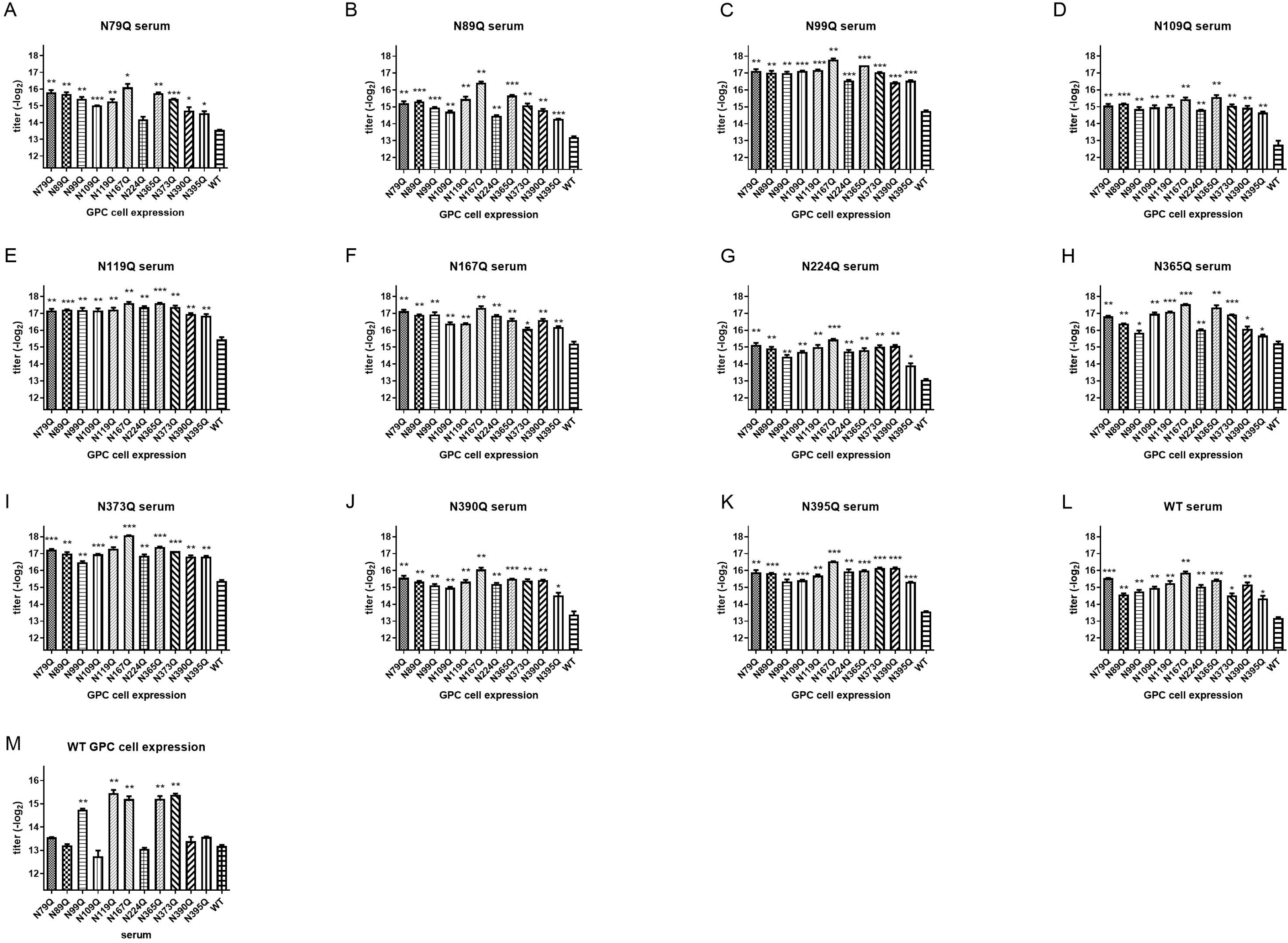
Antibody response to N-glycan deletion GPC showed higher titer to mutant GPC than to WT GPC. After three rounds of immunization, sera were collected and the titers were determined using cell-based ELISA. *P < 0.05, **P < 0.01, ***P < 0.001, compared to WT.

### Effects of N-glycosylation Modification on Antibody Neutralization

We then evaluated the neutralization ability of the serum against LASVpv infection (Fig. 6). Unfortunately, we found that the serum generated by both the WT GPC immunized group and each of the N-linked glycan deletion mutated variants showed minimal to no neutralizing effect to the LASVpv infection.

**Fig. 6.**
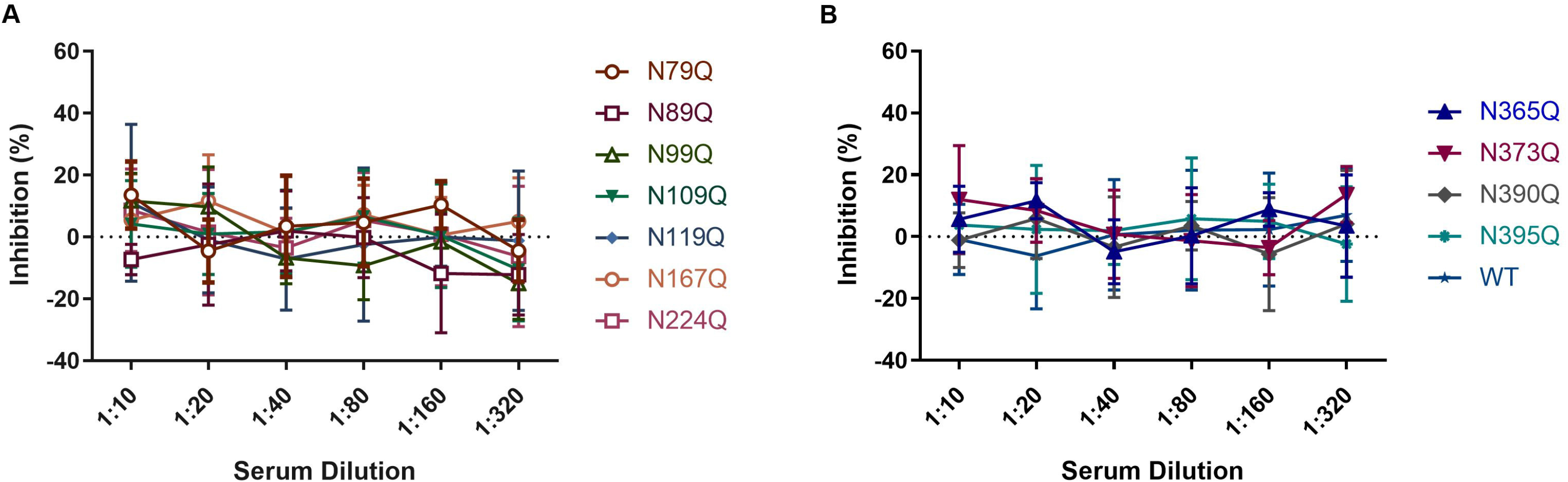
N-glycan deletions showed little effect in improving the neutralization ability of the antibody from sera against the seven individual glycan deletions in GP1 (A) and four in GP2 (B).

## Discussion

In a variety of virions such as influenza A virus, human immunodeficiency virus, Ebola virus, etc., a glycan shield on the glycoprotein plays critical roles in the host immune response. The LASV GPC harbors 11 N-linked glycans that almost completely encapsulate the GPC. Therefore, modification of these chains represents a possible strategy for identifying the LASV immune-escape mechanism, given that these chains might shield key epitopes on the protein surface from immune recognition. We hypothesized that removal of these chains could expose the epitopes, thereby promoting host recognition and triggering of the immune response. First, to investigate the role of the individual N-linked glycan in GPC expression and function, we found that using either stratagem to abolish the glycosylation motif, the deletion of the 2^nd^ (N89NS) or the 8^th^ (N365YS) glycan led to decreased GPC cleavage and pseudotype virus infectivity. These results were in line with previous reports that commented on the importance of these two glycans in LCMV GPC function (14, 15). As the 2^nd^ and 8^th^ glycans are absolutely conserved in *Mammarenavirus* GPC, it was supposed that both of the glycan chains play essential roles in GPC expression and function (1, 12). The 2^nd^ glycan was reported to interact with H92 in the prefusion conformation, and during endocytosis, the glycan chain was rotated and H92 was released, which could bind to the second receptor LAMP1 (1, 13). Similarly, the 2^nd^ glycan of the New World *Mammarenavirus* Machupo virus was reported to form a stacking interaction with F98, which was essential for receptor binding (12, 15). The 8^th^ glycan of LASV GPC was the first glycan in GP2. This glycan was reported to interact with Q232 and R235, shielding the fusion peptide at the tip of GP2, and thus contributing to the stability of the prefusion GPC (1).

It was recently reported that the 3^rd^ and 5^th^ glycan on LASV GPC shield the neutralizing epitopes of the virus (4, 8), and the 5^th^, 8^th^ and 9^th^ glycosylation motifs were reported to be located in the epitopes of GPC (8). Most recently, the 10^th^ and 11^th^ glycans were reported to occlude the conformational GPC-B epitope located at the stalk of GPC (16). Notably, evidence from human survivor and vaccine development studies have shown that adaptive immune protection in LASV infection is probably conferred mainly by a cell-mediated immune response, especially for Type I IFN (9, 17–23). It was supposed that deletion of the glycan chain would expose the epitope and thus increase the immunogenicity of GPC. We found that deletion of specific N-linked glycan residues on the GPC had no effect on the proportions of lymphocytes (CD3+, CD4+/CD3+, and CD8+/CD3+) but significantly increased proportions of effector lymphocytes (IFN-γ+/CD4+ and IFN-γ+/CD8+), especially following deletion of chains at the 1^st^, 3^rd^, 5^th^, and 6^th^ glycosylation sites. Additionally, we observed significant increases in the secretion of molecules involved in Th1 immune response (IL-2 and IFN-γ) following deletion of chains at the 3^rd^, 5^th^, 8^th^, 9^th^, and 10^th^ glycosylation sites, although this did not affect the secretion of Th2- and Th17-related cytokines. These results support the role of N-linked glycans in inhibiting host recognition and Th1 immune response.

Additionally, analysis of changes in antibody titers from the sera of mice immunized with GPC variants revealed general increases in titer following glycan removal, suggesting that the presence of these glycan chains promotes immune escape by shielding the antigenic epitope. Moreover, we verified that the antibodies generated by all GPC variants showed no neutralizing effect on the WT LASVpv. This indicated that the epitopes exposed by the deletion of N-linked glycans on the GPC did not generate neutralizing antibodies, suggesting that further investigation is required to identify these LASV epitopes.

In summary, we found that the N-sugar chains at the 3^rd^, 5^th^, 6^th^, 8^th^, and 9^th^ N-linked glycans likely shield epitopes on the LASV GPC that reduce host cellular and humoral immune responses. These sites can potentially be used as breakthrough points to develop effective therapeutic or prophylactic antibodies against Lassa fever.

## Materials and Methods

### Cells and Plasmids

HEK 293T, HeLa, and Vero cells were cultured in Dulbecco’s modified Eagle’s medium (HyClone, Logan, UT, USA) supplemented with 10% fetal bovine serum (FBS, Gibco, Grand Island, NY, USA). The pseudotype VSV bearing the GPC of LASV (strain Josiah, GenBank accession number HQ688673.1) as well as containing the *Renilla* luciferase (Rluc) reporter gene were generated as previously reported with the titer of 3 ×10^7^/mL (24).

To generate individual glycan deletion mutations, we introduced three amino acid substitutions (N→A, N→Q, and T/S→A) into the 11 N-glycosylation motifs using 33 pairs of primers (Table 1) (Synthesised by Sangon, Shanghai). The recombinant plasmids underwent PCR amplification to perform site-directed mutagenesis, after which the template was removed by *Dpn*I restriction digestion, and products were obtained via gel extraction, transformation, and monoclonal antibody identification.

**Table 1.**
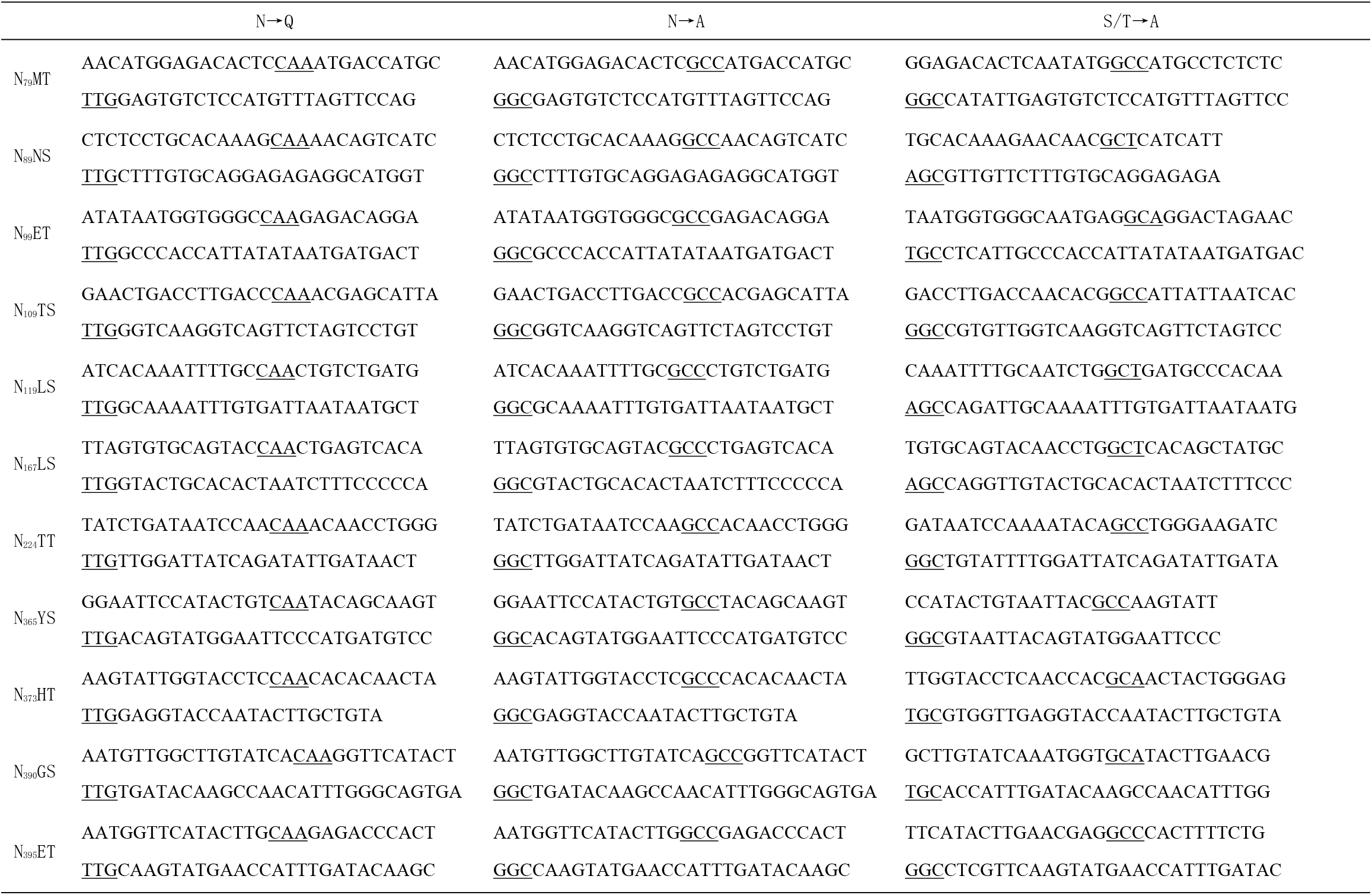
Primers for the mutagenesis.

### Mice

Specific pathogen-free (SPF) 6-week-old female BALB/c mice were maintained at the Laboratory Animal Center of Wuhan Institute of Virology, Chinese Academy of Sciences (CAS). All mouse studies were performed according to Regulations of the Administration of Affairs Concerning Experimental Animals in China (WIVA25201801), and the protocols were reviewed and approved by the Laboratory Animal Care and Use Committee at the Wuhan Institute of Virology, CAS. All mice were fed in independent ventilated cages (IVCs), and the IVCs were kept within an SPF barrier environment for experimental animals. The feed was sterilized via Co^60^ irradiation and water was sterilized using an autoclave.

### Immunization Strategy

Immunization was performed via intramuscular injection using 40 μg of each respective plasmid in the medial thigh of each mouse while avoiding blood vessels. To improve immunogenicity, mice were shocked with an Electro Square Porator (BTX, ECM830) using the cross method with the injection hole as the center. Six mice in each group were immunized three times over a 2-week interval, and at 10 days after the final immunization, mice were euthanized via decapitation; eyeball enucleation was conducted for blood collection.

### Separation of Mouse Spleen Lymphocytes

Lymphocyte separation was performed using an EZ-Sep kit (Dakewe Biotech Co., Ltd., Beijing, China) according to manufacturer’s instructions. Mouse spleen were soaked in Roswell Park Memorial Institute (RPMI) 1640 medium. EZ-Sep separation solution (3-4 mL) was then added to a sterile 3 cm culture dish, over which a nylon mesh was fixed with hemostatic forceps. The spleen were placed onto the mesh for grinding and grinding solution was rapidly transferred to a 15 mL centrifuge tube along with ~500 μL of serum-free RPMI 1640 along the tube wall; the solution was centrifuged at 800 *g* at 25 °C for 30 min. The lymphocyte layer was transferred to a new 15 mL centrifuge tube, followed by the addition of 10 mL serum-free RPMI 1640 medium and centrifugation at 250 *g* for 10 min. The supernatant was then carefully removed, the cells were resuspended with 500 μL RPMI 1640 medium containing 10% FBS, and 10 μL was used for 10-fold serial dilutions for cell counting. The solution was diluted to a cell density of 2×10^7^ cells/mL, transferred to a 96-well U-shaped-bottom plate at 100 μL/well, and cultured in a cell incubator with 5% CO_2_ at 37 °C.

### Detection of CD4+ and CD8+ T Cells and Cytokines

The culture system used to stimulate spleen lymphocytes involved the application of a stimulator and co-stimulator [(anti-CD28) + Golgi blocker (BFA) + spleen lymphocytes (2 × 10^6^ cells)]. Negative control, positive control, and experimental groups underwent stimulation with phosphate-buffered saline (PBS), phorbol myristate acetate/ionomycin, and polypeptide (Table 2) (Synthesised by Bankpeptide, Heifei) (25), respectively. After 4.5 h in culture, the solution was centrifuged at 800 *g* for 3 min; 50 μL TruStain FcX™ (mouse anti-CD16/32; BioLegend, San Dieg, CA, USA) was added to reduce non-specific fluorescent staining, incubated at 4 °C for 10 min, and centrifuged at 800 *g* for 3 min. PBS solution was then used to dilute fluorescently labeled antibodies (PE-conjugated rat anti-mouse CD8a, FITC-conjugated rat anti-mouse CD4, PE–Cy™7-conjugated hamster anti-mouse CD3e), and 7-AAD viability staining solution (BioLegend) by 1:200. They were then added (100 μL/well) for staining at 4 °C for 20 min in the dark. After washing, the cells were fixed, permeabilized, and subjected to intracellular staining. An allophycocyanin-conjugated rat anti-mouse IFN-γ antibody (1:200) was then added and incubated at 4 °C for 30 min in the dark. The cells were filtered through a 200 μm nylon mesh before being loaded onto the flow cytometer (BD FACSAria III). Cytokines were detected using a cytometric bead array kit (BD Biosciences, Franklin Lakes, NJ, USA) according to manufacturer’s instructions.

**Table 2.** Simulation peptides (predicted by www.iedb.org)

|  | location | sequencing | purity |
| --- | --- | --- | --- |
| P1 | 277-285 | GYCLTRWML | 95% |
| P2 | 128-136 | LYDHALMSI | 95% |
| P3 | 65-73 | VYELQTLEL | 95% |
| P4 | 315-323 | LFDFNKQAI | 95% |
| P5 | 156-164 | DFNGGKISV | 95% |
| P6 | 322-336 | AIQRLKAEAQMSIQL | 95% |

### Antibody Titration

The sera were collected from each immunized mouse 10 d after the last immunization to determine specific IgG using cell-based ELISA. The serum was diluted 50-fold, followed by separation into eight gradient dilutions at 1:4 ratios. HeLa cells were transfected with the individual glycan deletions with WT and control plasmids serving as antigens, which were blocked, washed, and incubated with serums, followed by detection with HRP-conjugated AffiniPure Goat Anti-Mouse IgG (Proteintech, Wuhan, China).

### Antibody Neutralization

Serum was diluted 10-fold with FBS-free medium and then separated into six gradients at 1:2 ratios. Forty microliters of the diluted serum were mixed with 10 μL LASVpv at 37 °C for 1 h. The mixture was added to Vero cells for 1 h incubation. Neutralization activities were determined 24 hours later using the Rluc assay system (Promega, Madison, WI, USA).

## ACKNOWLEDGEMENTS

We thank the Center for Instrumental Analysis and Metrology, Core Facility and Technical Support, and Center for Animal Experiment, Wuhan Institute of Virology, for providing technical assistance. We would like to thank Editage (www.editage.cn) for English language editing

This work was supported by the National Key Research and Development Program of China (2018YFA0507204), the National Natural Sciences Foundation of China (31670165), Wuhan National Biosafety Laboratory, Chinese Academy of Sciences Advanced Customer Cultivation Project (2019ACCP-MS03), the Open Research Fund Program of the State Key Laboratory of Virology of China (2018IOV001).

## REFERENCES

1. Hastie KM, Zandonatti MA, Kleinfelter LM, Heinrich ML, Rowland MM, Chandran K, Branco LM, Robinson JE, Garry RF, Saphire EO. 2017. Structural basis for antibody-mediated neutralization of Lassa virus. Science 356:923–928.

2. Eichler R, Lenz O, Garten W, Strecker T. 2006. The role of single N-glycans in proteolytic processing and cell surface transport of the Lassa virus glycoprotein GP-C. Virol J 3:41.

3. Watanabe Y, Raghwani J, Allen JD, Seabright GE, Li S, Moser F, Huiskonen JT, Strecker T, Bowden TA, Crispin M. 2018. Structure of the Lassa virus glycan shield provides a model for immunological resistance. Proc Natl Acad Sci U S A doi:10.1073/pnas.1803990115.

4. Sommerstein R, Flatz L, Remy MM, Malinge P, Magistrelli G, Fischer N, Sahin M, Bergthaler A, Igonet S, Ter Meulen J, Rigo D, Meda P, Rabah N, Coutard B, Bowden TA, Lambert PH, Siegrist CA, Pinschewer DD. 2015. Arenavirus Glycan Shield Promotes Neutralizing Antibody Evasion and Protracted Infection. PLoS Pathog 11:e1005276.

5. Johnson KM, McCormick JB, Webb PA, Smith ES, Elliott LH, King IJ. 1987. Clinical virology of Lassa fever in hospitalized patients. J Infect Dis 155:456–64.

6. Gunther S, Emmerich P, Laue T, Kuhle O, Asper M, Jung A, Grewing T, ter Meulen J, Schmitz H. 2000. Imported lassa fever in Germany: molecular characterization of a new lassa virus strain. Emerg Infect Dis 6:466–76.

7. McCormick JB, King IJ, Webb PA, Scribner CL, Craven RB, Johnson KM, Elliott LH, Belmont-Williams R. 1986. Lassa fever. Effective therapy with ribavirin. N Engl J Med 314:20–6.

8. Robinson JE, Hastie KM, Cross RW, Yenni RE, Elliott DH, Rouelle JA, Kannadka CB, Smira AA, Garry CE, Bradley BT, Yu H, Shaffer JG, Boisen ML, Hartnett JN, Zandonatti MA, Rowland MM, Heinrich ML, Martinez-Sobrido L, Cheng B, de la Torre JC, Andersen KG, Goba A, Momoh M, Fullah M, Gbakie M, Kanneh L, Koroma VJ, Fonnie R, Jalloh SC, Kargbo B, Vandi MA, Gbetuwa M, Ikponmwosa O, Asogun DA, Okokhere PO, Follarin OA, Schieffelin JS, Pitts KR, Geisbert JB, Kulakoski PC, Wilson RB, Happi CT, Sabeti PC, Gevao SM, Khan SH, Grant DS, Geisbert TW, Saphire EO, Branco LM, Garry RF. 2016. Most neutralizing human monoclonal antibodies target novel epitopes requiring both Lassa virus glycoprotein subunits. Nat Commun 7:11544.

9. Meulen J, Badusche M, Satoguina J, Strecker T, Lenz O, Loeliger C, Sakho M, Koulemou K, Koivogui L, Hoerauf A. 2004. Old and New World arenaviruses share a highly conserved epitope in the fusion domain of the glycoprotein 2, which is recognized by Lassa virus-specific human CD4+ T-cell clones. Virology 321:134–43.

10. ter Meulen J, Badusche M, Kuhnt K, Doetze A, Satoguina J, Marti T, Loeliger C, Koulemou K, Koivogui L, Schmitz H, Fleischer B, Hoerauf A. 2000. Characterization of human CD4(+) T-cell clones recognizing conserved and variable epitopes of the Lassa virus nucleoprotein. J Virol 74:2186–92.

11. Bowden TA, Crispin M, Graham SC, Harvey DJ, Grimes JM, Jones EY, Stuart DI. 2009. Unusual molecular architecture of the machupo virus attachment glycoprotein. J Virol 83:8259–65.

12. Abraham J, Corbett KD, Farzan M, Choe H, Harrison SC. 2010. Structural basis for receptor recognition by New World hemorrhagic fever arenaviruses. Nat Struct Mol Biol 17:438–44.

13. Cohen-Dvashi H, Cohen N, Israeli H, Diskin R. 2015. Molecular Mechanism for LAMP1 Recognition by Lassa Virus. J Virol 89:7584–92.

14. Bonhomme CJ, Capul AA, Lauron EJ, Bederka LH, Knopp KA, Buchmeier MJ. 2011. Glycosylation modulates arenavirus glycoprotein expression and function. Virology 409:223–33.

15. Bonhomme CJ, Knopp KA, Bederka LH, Angelini MM, Buchmeier MJ. 2013. LCMV glycosylation modulates viral fitness and cell tropism. PLoS One 8:e53273.

16. Hastie KM, Cross RW, Harkins SS, Zandonatti MA, Koval AP, Heinrich ML, Rowland MM, Robinson JE, Geisbert TW, Garry RF, Branco LM, Saphire EO. 2019. Convergent Structures Illuminate Features for Germline Antibody Binding and Pan-Lassa Virus Neutralization. Cell 178:1004–1015 e14.

17. Ibukun FI. 2020. Inter-Lineage Variation of Lassa Virus Glycoprotein Epitopes: A Challenge to Lassa Virus Vaccine Development. Viruses 12.

18. Baize S, Marianneau P, Loth P, Reynard S, Journeaux A, Chevallier M, Tordo N, Deubel V, Contamin H. 2009. Early and strong immune responses are associated with control of viral replication and recovery in lassa virus-infected cynomolgus monkeys. J Virol 83:5890–903.

19. Warner BM, Safronetz D, Stein DR. 2018. Current research for a vaccine against Lassa hemorrhagic fever virus. Drug Des Devel Ther 12:2519–2527.

20. Mire CE, Cross RW, Geisbert JB, Borisevich V, Agans KN, Deer DJ, Heinrich ML, Rowland MM, Goba A, Momoh M, Boisen ML, Grant DS, Fullah M, Khan SH, Fenton KA, Robinson JE, Branco LM, Garry RF, Geisbert TW. 2017. Human-monoclonal-antibody therapy protects nonhuman primates against advanced Lassa fever. Nat Med doi:10.1038/nm.4396.

21. Geisbert TW, Jones S, Fritz EA, Shurtleff AC, Geisbert JB, Liebscher R, Grolla A, Stroher U, Fernando L, Daddario KM, Guttieri MC, Mothe BR, Larsen T, Hensley LE, Jahrling PB, Feldmann H. 2005. Development of a new vaccine for the prevention of Lassa fever. PLoS Med 2:e183.

22. Botten J, Alexander J, Pasquetto V, Sidney J, Barrowman P, Ting J, Peters B, Southwood S, Stewart B, Rodriguez-Carreno MP, Mothe B, Whitton JL, Sette A, Buchmeier MJ. 2006. Identification of protective Lassa virus epitopes that are restricted by HLA-A2. J Virol 80:8351–61.

23. Fisher-Hoch SP, Hutwagner L, Brown B, McCormick JB. 2000. Effective vaccine for lassa fever. J Virol 74:6777–83.

24. Wang P, Liu Y, Zhang G, Wang S, Guo J, Cao J, Jia X, Zhang L, Xiao G, Wang W. 2018. Screening and Identification of Lassa Virus Entry Inhibitors from an FDA-Approved Drugs Library. J Virol 92:e00954–18.

25. Vita R, Mahajan S, Overton JA, Dhanda SK, Martini S, Cantrell JR, Wheeler DK, Sette A, Peters B. 2019. The Immune Epitope Database (IEDB): 2018 update. Nucleic Acids Res 47:D339–D343.

